# The Specificity of IQGAP1 Toward the PI3K-Akt Pathway is Dependent on the IQ3 motif

**DOI:** 10.1101/250936

**Authors:** Mo Chen, Tianmu Wen, Suyong Choi, Narendra Thapa, Oisun Jung, Paul F. Lambert, Alan C. Rapraeger, Richard A. Anderson

## Abstract

Epidermal growth factor receptor (EGFR) and its downstream phosphatidylinositol 3-kinase (PI3K) pathway are commonly deregulated in many cancers including head and neck cancer (HNC). Recently, we have shown that the IQ motif-containing GTPase-activating protein 1 (IQGAP1) provides a molecular platform to scaffold all the components from the PI3K-Akt pathway and results in the sequential generation of phosphatidylinositol-3,4,5-triphosphate (PI3,4,5P_3_). This makes the IQGAP1-PI3K scaffold a promising therapeutic target. In addition to the PI3K-Akt pathway, IQGAP1 also scaffolds the Ras-ERK pathway. To identify an IQGAP1 mutant that specifically loses IQGAP1-PI3K signaling but not other functions, we have focused on the IQ3 motif since this region binds with both the PIPK1α and PI3K enzymes and a short peptide derived from this sequence blocks binding and PI3K signaling. An IQ3 deletion mutant (ΔIQ3) in IQGAP1 was functionally compared with wild-type (WT) IQGAP1. We found that the IQ3 domain specifically mediates the PI3K-Akt pathway but does not regulate the Ras-ERK pathway. The IQ3 deletion mutant lost interactions with PI3K-Akt components but retained its binding of ERK pathway components and cell surface receptors, EGFR and integrins. In addition, the IQ3 deletion mutant lost regulation of cell migration. Consistently, the IQ3 motif derived peptide blocked Akt activation and led to the blockage of invasion mediated by EGFR organized into an integrin complex by syndecan-4. Taken together, this work has defined the IQGAP1 IQ3 motif as a specific target sequence for the scaffolding of the PI3K-AKT pathway.

## Introduction

Epidermal growth factor receptor (EGFR) and its downstream phosphatidylinositol 3-kinase (PI3K) pathway are commonly deregulated in HNC, making them promising therapeutic targets^1–3^. We have recently discovered that the multi-domain scaffolding IQ motif-containing GTPase-activating protein 1 (IQGAP1) provides a novel molecular platform for the assembly of PI4P-, PI4,5P_2_- and PI3,4,5P_3_-generating enzymes (PI4KIIIα, PIPKIα and PI3K) into proximity for concerted and efficient generation of the PI3,4,5P_3_ lipid messenger^4^. The generated PI3,4,5P_3_ recruits PDK1/Akt kinases to the IQGAP1 complex and activates them. This finding suggests that IQGAP1 is a novel therapeutic target for cancer treatment.

IQGAP1, also known as p195, is a multidomain protein first discovered in 1994^7^. It belongs to the IQGAP family, together with IQGAP2 and IQGAP3. The domains in IQGAP1 are, starting from the N-terminus, the calponin homology domain (CHD), the poly-proline protein-protein domain (WW), the 4 IQ domains and the RasGAP-related domain (GRD) (Figure 1a). In cells, IQGAP1 functions as a scaffold protein, playing a key role in both the PI3K-Akt and Ras-ERK pathways^8^. Crucial components of these pathways are assembled by IQGAP1. This includes the full PI3K-Akt pathway components (PI4KIIIα, PIPKIα, Ras, PI3K, PDK1, and AKT) and the Ras-ERK pathway components (Ras, Raf, MEK, and ERK) are closely linked together by IQGAP1^9^. The Ras-ERK and PI3K-Akt pathways are key mechanisms for controlling cell growth, survival, metabolism, and motility^10^. In response to agonist activation, these two pathways crosstalk to regulate downstream cellular signaling and also each other. IQGAP1 has been shown to mirror these functions and is consistent with scaffolding both pathways. Enhanced activation of one pathway that occurs by inhibition of the other has been considered as a mechanism for chemoresistance in cancer treatment^11^. For example, FOXO1, the downstream target of Akt, also binds to IQGAP1 and phosphorylated FOXO1 inhibits the Ras-ERK pathway^12^. The tight association between the PI3K-Akt and the Ras-ERK pathways coupled by IQGAP1 illustrates the importance of identifying specific sequences on IQGAP1 that selectively control the PI3K-Akt or Ras-ERK pathways.

**Figure 1.**
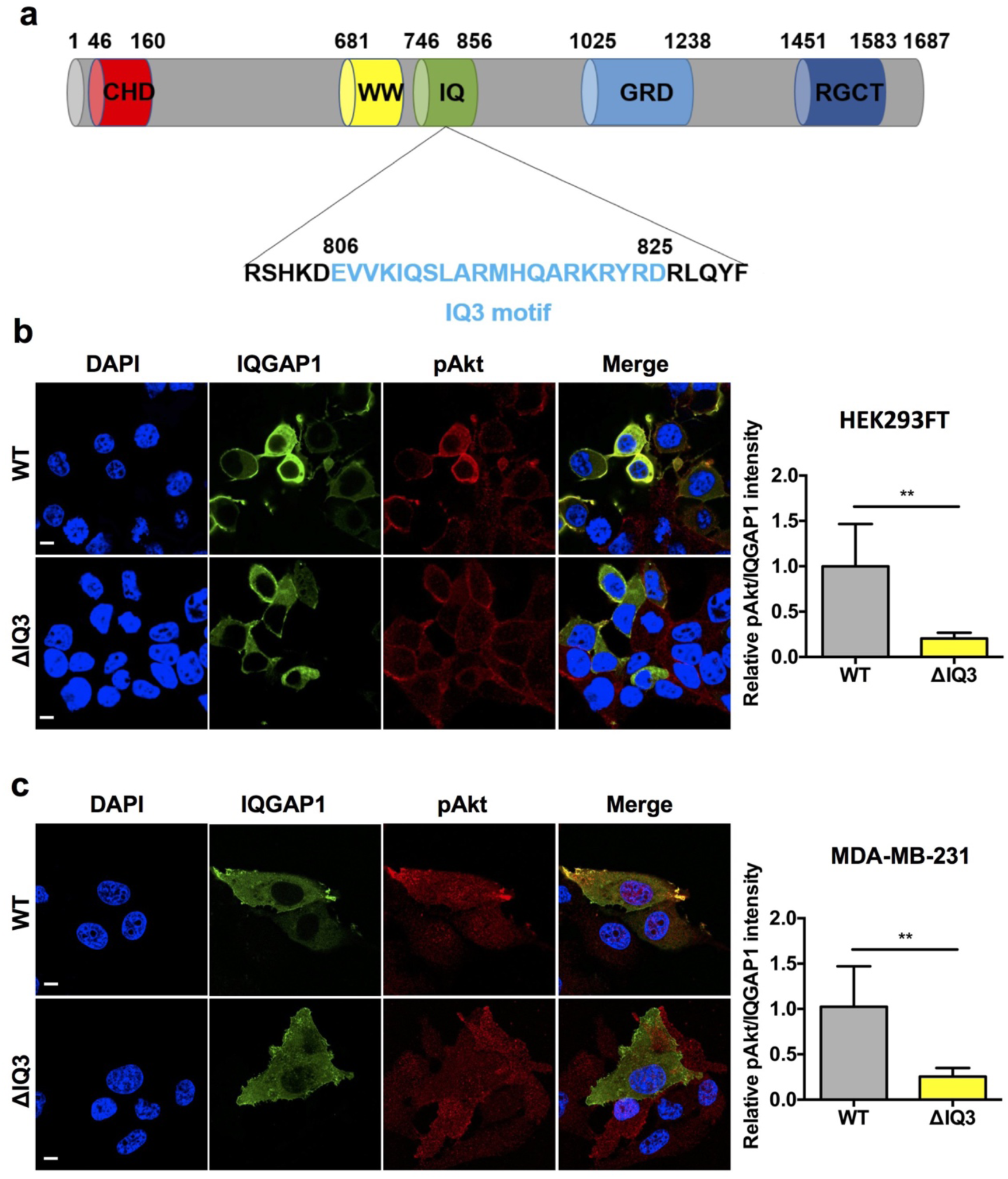
The IQ3 motif in IQGAP1 mediates the PI3K-Akt pathway. (**a**) Domain model of IQGAP1. (**b, c**) Deletion of the IQ3 motif in IQGAP1 blocked the inducible activation of the PI3K-Akt pathway. HEK293FT and MDA-MB-231 cells post 24 h of transient transfection with GFP-tagged IQGAP1^WT^ or GFP-tagged IQGAP1^ΔIQ3^ were starved for 24 h and then treated with 10 ng/ml EGF for 15 min. The cells were fixed by 4% PFA in PBS and processed for immunofluorescent staining of pAkt. The nuclei were counterstained with DAPI. The images were taken by Leica SP8 confocal microscope and quantified by ImageJ. The signal intensity of pAkt (Red) channel divided by the signal intensity of IQGAP1 (Green) channel was used as the indicator for the effect of IQGAP1 WT and ΔIQ3 mutant on the PI3K-Akt pathway. \*\**P*<0.01, n=10. Error bars denote SD. Scale bar, 5 μm.

The scaffolding of PI3K-Akt pathway components by IQGAP1 positions the phosphoinositide kinases into functional proximity for efficient generation of PIP_3_ and Akt activation^4^. This indicates that each of the kinases binds to a dedicated sequence of IQGAP1. A key region is the IQ3 domain (Figure 1a). It has been shown that the IQ3 domain selectively binds to both PIPKIα and the PI3K regulatory subunit, p85α and PIPKIα and p85α are essential components of PI3K-Akt pathway^4^. However, whether IQ3 motif mediates the interaction with PIPKIα and p85α in the context of full-length IQGAP1 is still unknown and whether the IQ3 sequence is specific for the PI3K pathway compared to the ERK pathway has not been defined.

Here, we set out to determine if the IQ3 domain in IQGAP1 is a specific binding site for PI3K-Akt pathway components that is separated from the upstream activating receptors of the PI3K-Akt or Ras-ERK pathways. For this, we constructed an IQ3 deletion mutant (ΔIQ3) and studied its impact on the PI3K-Akt and Ras-ERK pathways.

The results show that the IQ3 domain is required for PI3K-Akt pathway signaling but does not regulate the Ras-ERK pathway or IQGAP1 interaction with receptors. Consistently, the IQ3 deletion mutant lost the interaction with the PI3K-Akt components but retained binding to both ERK pathway components and cell surface receptors,including EGFR, integrin a3b1, and integrin a6b4. In addition, the IQ3 deletion mutant lost regulation of cell migration and an IQ3 motif-derived peptide both reduced Akt activation and blocked HNC cell invasion that is mediated by EGFR and the a3b1 and integrin a6b4 when organized as a receptor complex by syndecan-4 (the Sdc4:EGFR:ITG complex). Taken together, this work has demonstrated that the IQ3 motif is a specific target in IQGAP1 for the PI3K-Akt pathway.

## Results

### IQ3 motif of IQGAP1 mediates the activation of Akt pathway

The Myc-tagged and GFP-tagged WT IQGAP1 has been shown to fully rescue IQGAP1 deletion phenotypes^4,13,14^. Therefore, in order to determine the contribution of IQ3 domain to the activation of the PI3K-Akt pathway, human embryonic kidney cells (HEK293FT) or human breast carcinoma cells (MDA-MB-231) transiently transfected with Myc-tagged or GFP-tagged WT or IQ3 motif deleted IQGAP1 (IQGAP1^ΔIQ3^) expression plasmids were utilized (Figure 1a). Whereas WT IQGAP1 stimulated Akt phosphorylation in serum-starved HEK293FT cells upon EGF stimulation and the level of Akt phosphorylation corresponded with the expression level of WT IQGAP1, the IQ3 deletion mutant of IQGAP1 failed to activate Akt (Figure 1b). There was no difference in the Akt activation between the untransfected cells and IQGAP1^ΔIQ3^ transfected cells (Figure 1b). Through quantification of the relative pAkt intensity in the GFP positive cells, there was a considerable loss of activation between the WT and IQ3 deletion mutant. These results were further confirmed by using MDA-MB-231 cells (Figure 1c), indicating that the IQ3 sequence in IQGAP1 is required for Akt activation.

### Deletion of IQ3 motif in IQGAP1 does not affect Ras-ERK signaling

In addition to PI3K-Akt components, IQGAP1 harbors binding sites for Ras-ERK signaling pathway. Therefore, to assess the specificity of the IQ3 domain in the regulation of the PI3K-Akt pathway, independent of the Ras-ERK pathway, ERK phosphorylation was analyzed in HEK293FT and MDA-MB-231 cells transiently expressing GFP-tagged IQGAP1^WT^ and IQGAP1^ΔIQ3^. Immunostaining followed by confocal microscopy was performed to detect changes in phosphorylation of ERK, an indicator for the activation of the Ras-ERK pathway. Results showed that ERK phosphorylation was induced to a similar extent, irrespective of WT or IQ3 deletion mutant of IQGAP1 was overexpressed (Figure 2a), and the amount of ERK phosphorylation was correlated with the level of expression of WT or IQGAP1^ΔIQ3^. Additionally, no difference between WT and IQ3 deletion mutant of IQGAP1 on the ERK activation was detected. In contrast, MDA-MB-231 cells have been shown to exhibit higher endogenous levels of ERK activation due to dominant active Ras. As a consequence, the overexpression of WT IQGAP1 or IQGAP1^ΔIQ3^ in these cells did not enhance the ERK activation than the untransfected cells (Figure 2b). Thus, the absence of detectable difference in the effect of WT or IQGAP1^ΔIQ3^ on ERK phosphorylation suggests that the IQ3 motif within IQGAP1 does not affect ERK pathway.

**Figure 2.**
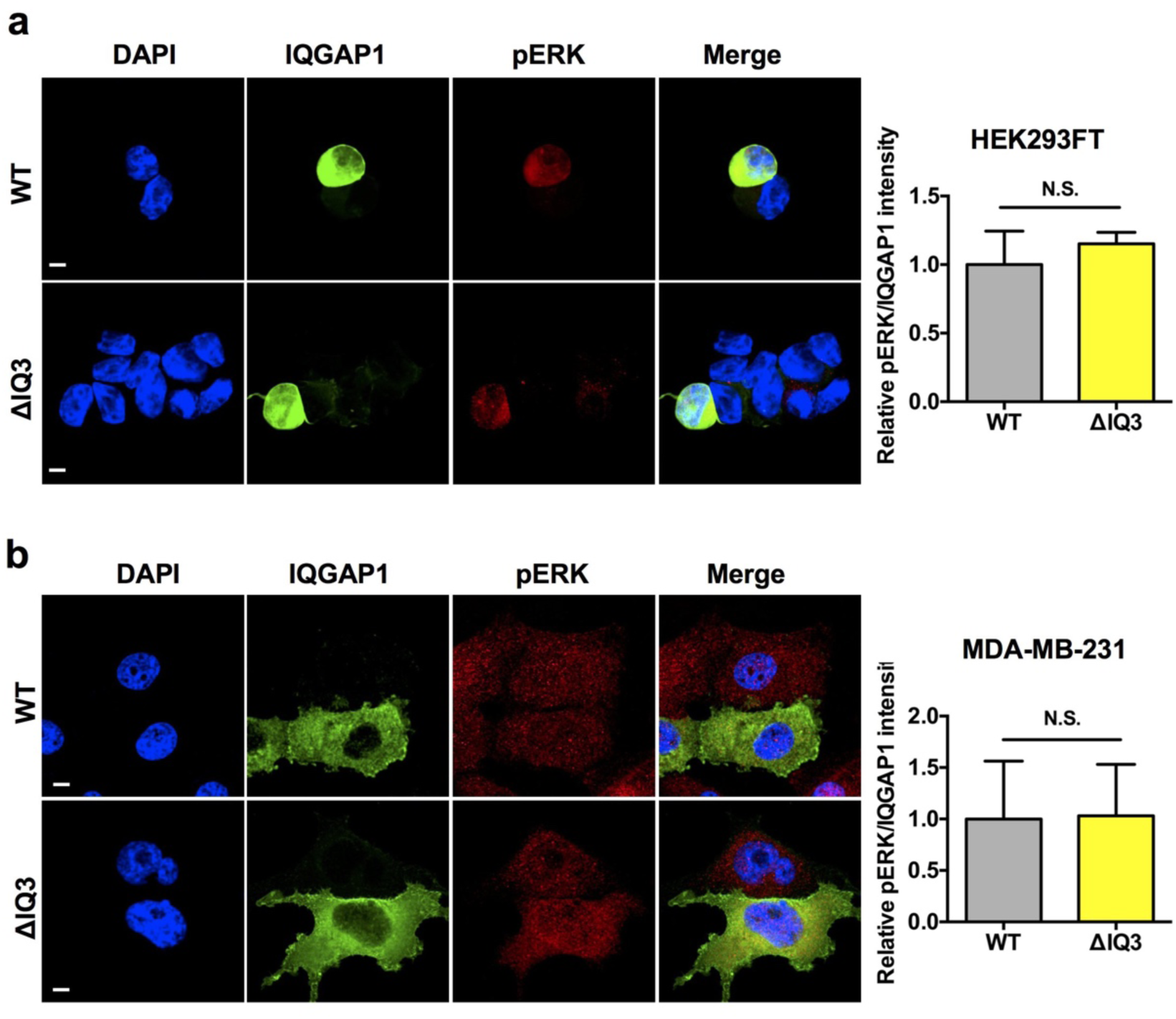
Deletion of the IQ3 motif in IQGAP1 has no impact on the ERK pathway. (a, b) Deletion of the IQ3 motif in IQGAP1 has no influence on the ERK activation. HEK293FT and MDA-MB-231 cells post 24 h of transient transfection with GFP-tagged IQGAP1^WT^ or GFPtagged IQGAP1^ΔIQ3^ were starved for 24 h and then treated with 10 ng/ml EGF for 15 min. The cells were fixed by 4% PFA in PBS and processed for immunofluorescent staining of pERK. The nuclei were counterstained with DAPI. The images were taken by Leica SP8 confocal microscope and quantified by ImageJ. The signal intensity of pERK (Red) channel divided by the signal intensity of IQGAP1 (Green) channel was used as the indicator for the effect of IQGAP1 WT and ΔIQ3 mutant on the ERK pathway. n=10. Error bars denote SD. Scale bar, 5 μm.

### IQ3 motif in IQGAP1 selectively activates the PI3K-Akt pathway

To confirm the confocal microscopy results, Myc-tagged IQGAP1^WT^ and IQGAP1^ΔIQ3^ expressing HEK293FT cells were subjected similar treatments as in microscopic analysis. Immunoblotting was performed to analyze the phosphorylation level of Akt and ERK. ERK phosphorylation was induced upon EGF stimulation and was further enhanced by the overexpressing of both WT and the IQ3 deletion mutant of IQGAP1 (Figure 3a-b). Whereas Akt phosphorylation was upregulated only in cells expressing WT-IQGAP1 but not IQGAP1^ΔIQ3^, activated Akt levels remained unchanged in cells expressing IQGAP1^ΔIQ3^as in non-transfected control cells (Figure 3c). Taken together, these data suggest that IQ3 motif is required for the activation of Akt pathway independent of Ras-ERK signaling.

**Figure 3.**
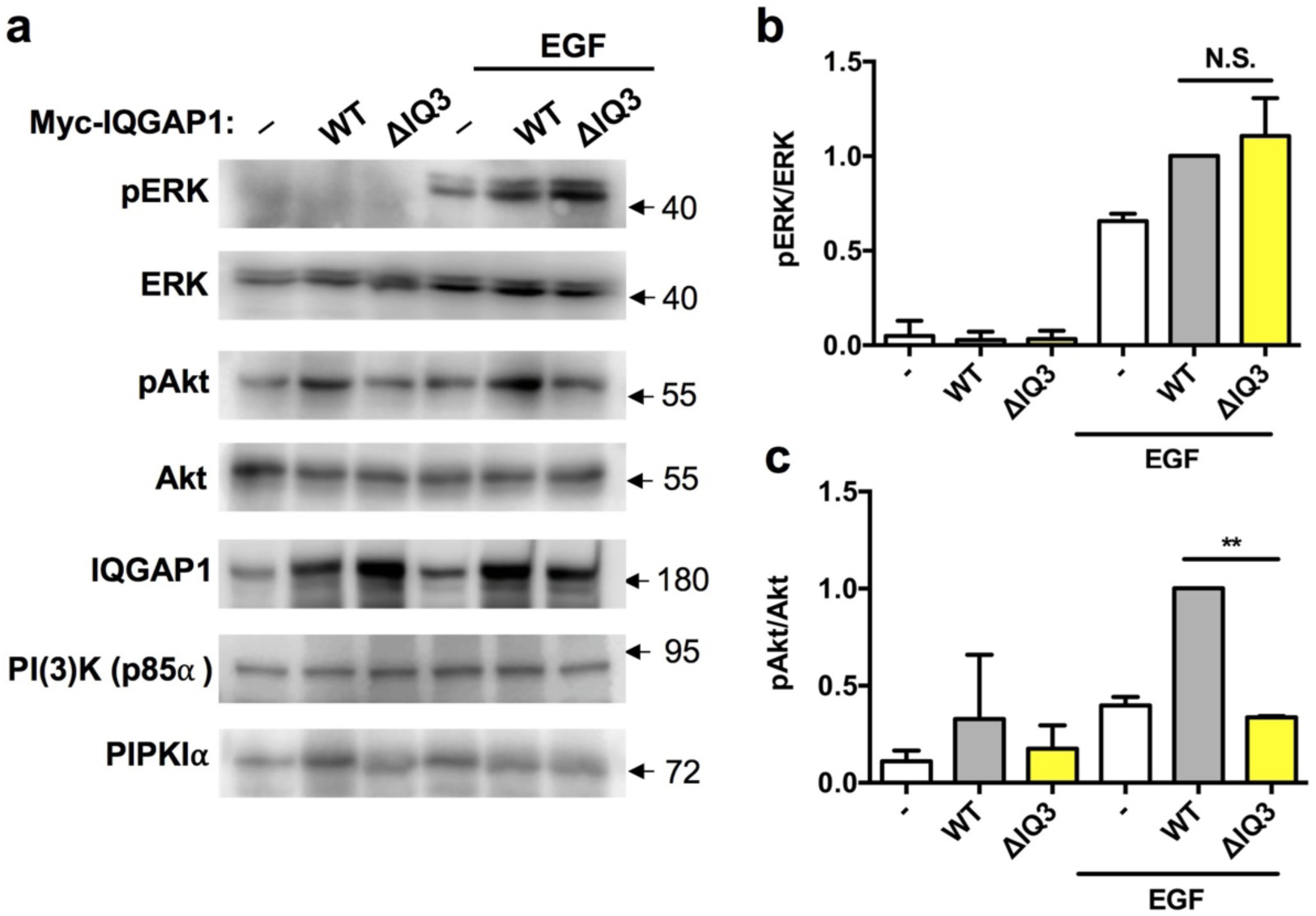
The IQ3 motif in IQGAP1 specifically mediates the PI3K-Akt pathway in HEK293FT cells. (**a**) EGF stimulated pAkt and pERK were measured in HEK293FT cells. HEK293FT cells post 24 h of transient transfection with Myc-tagged IQGAP1^WT^ or Myc-tagged IQGAP1ΔIQ3 were starved for 24 h and then treated with 10 ng/ml EGF for 15 min as indicated. The whole cell lysates were collected and processed for WB. (**b-c**) Quantification of relative pERK and pAkt level in **a**. ⋆⋆P<0.01, n=3. Error bars denote SD.

### IQ3 motif in IQGAP1 physically interacts with components of PI3K-Akt pathway

To further validate the role of IQ3 motif in selective activation of PI3K-Akt pathway, the interaction between IQ3 motif and components of PI3K-Akt as well as Ras-ERK signaling pathways were investigated. Cells expressing Myc-tagged IQGAP1^WT^ and IQGAP1^ΔIQ3^ were subjected to immunoprecipitation using anti-Myc antibodies followed by immunoblotting to identify and quantify the associated proteins. The results showed that EGF stimulation enhanced the interaction between WT IQGAP1 and all of the examined components of PI3K-Akt and Ras-ERK pathway, including, total ERK, pERK, total Akt, pAkt, PDK1, p85, and PIPK1α (Figure 4a). The IQGAP1^ΔIQ3^ was able to interact with ERK (Figure 4b) and pERK (Figure 4c). However, there was a significant reduction in the binding of the IQGAP1^ΔIQ3^ to Akt (Figure 4d), pAkt (Figure 4e), PDK1 (Figure 4f), p85α (Figure 4g) and PIPK1α (Figure 4h) compared to WT IQGAP1. The binding between PIPK1α and IQGAP1 was completely lost when the IQ3 motif was deleted while the binding between p85α and IQGAP1 was partially reduced. Consistently, there was also a complete loss of pAKT. These results are consistent with previous studies showing that IQ3 motif is the binding site for both PIPK1α and p85α^4^. However, p85α also binds with the WW domain consistent with its partial reduction in binding. These results indicate that the IQ3 sequence in IQGAP1 selectively interacts with PI3K-Akt pathway components.

**Figure 4.**
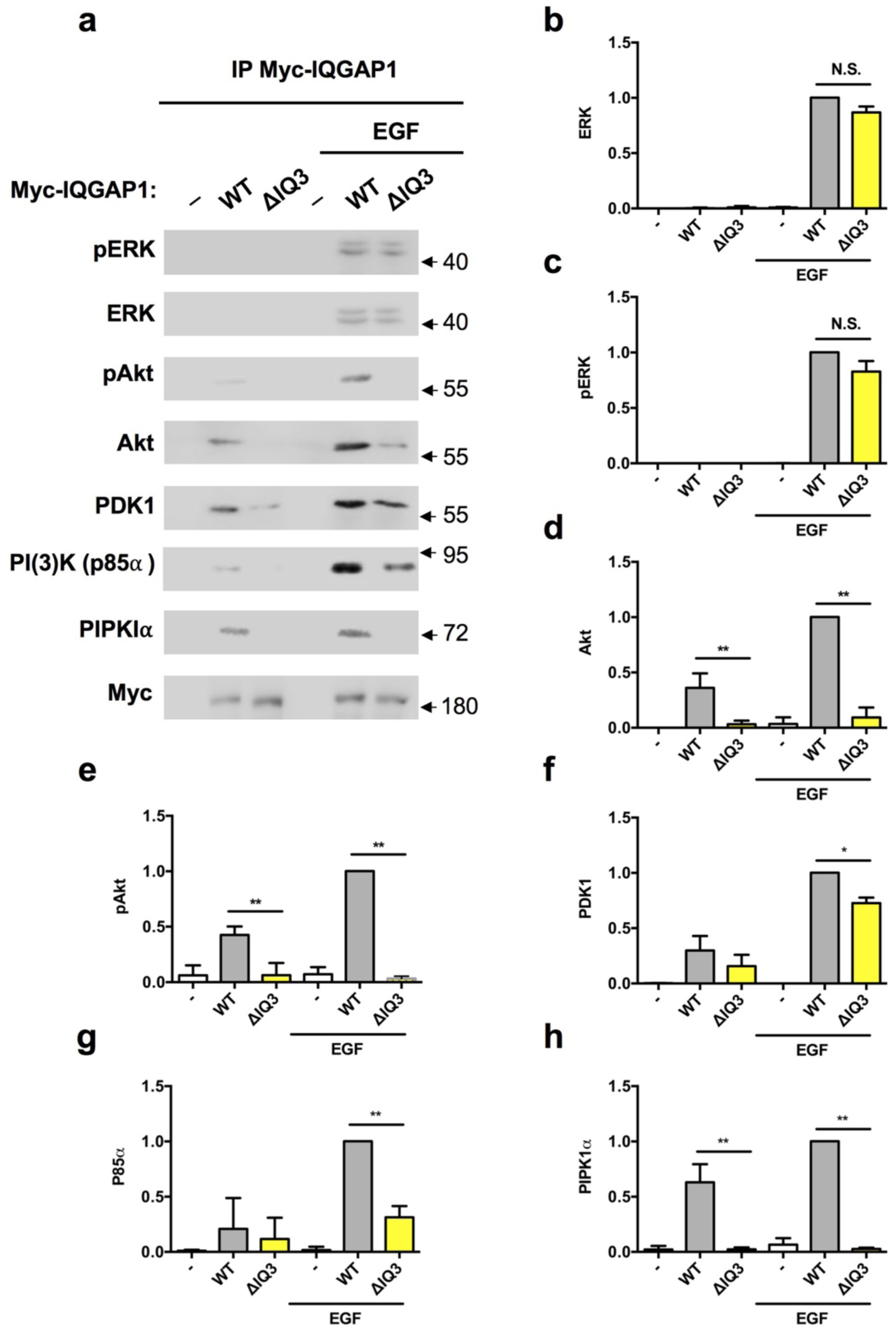
IQ3 motif specifically mediates the interactions between IQGAP1 and PI3K-Akt pathway components. (**a**) IQGAP1 WT and ΔIQ3 mutant associated ERK/PI3K-Akt pathway components were analyzed by immunoprecipitation. HEK293FT cells post 24 h of transient transfection with Myc-tagged IQGAP1^WT^ or Myc-tagged IQGAP1^ΔIQ3^ were starved for 24 h and then treated with 10 ng/ml EGF for 15 min as indicated. The whole cell lysates were collected and processed for immunoprecipitation using anti-Myc antibody conjugated Protein A/G agarose beads. (**b-h**)
Quantification of immunoblots in **a**. ⋆*P*<0.05, ⋆⋆P<0.01, n=3. Error bars denote SD.

### IQ3 deletion in IQGAP1 retains the binding to cell surface receptors

IQGAP1 functions to scaffold PI3K and ERK upon agonist (e.g. EGF) stimulation and other extracellular stimuli through cell surface receptors. Further, EGFR, integrin α3β1, and integrin α6β4 have been reported to be organized into a complex by syndecan-4 to exert downstream signaling and promote cell invasion^15–17^ (the Sdc4:EGFR:ITG complex). Therefore, we assayed the binding between WT/IQ3 deletion mutant of IQGAP1 and these cell surface receptors. Integrin α3 subunit only forms a heterodimer with integrin β1 subunit and integrin β4 subunit only forms a heterodimer with integrin α6 subunit^18^. Therefore, immunoblotting of integrin α3 subunit is a representative of integrin α3β1 complex. Similarly, immunoblotting of integrin β4 subunit is sufficient to indicate integrin α6β4 complex. The results show that WT and IQ3 deletion mutant of IQGAP1 bind to EGFR, integrin α3β1, and integrin α6β4 indistinguishably (Figure 5).

**Figure 5.**
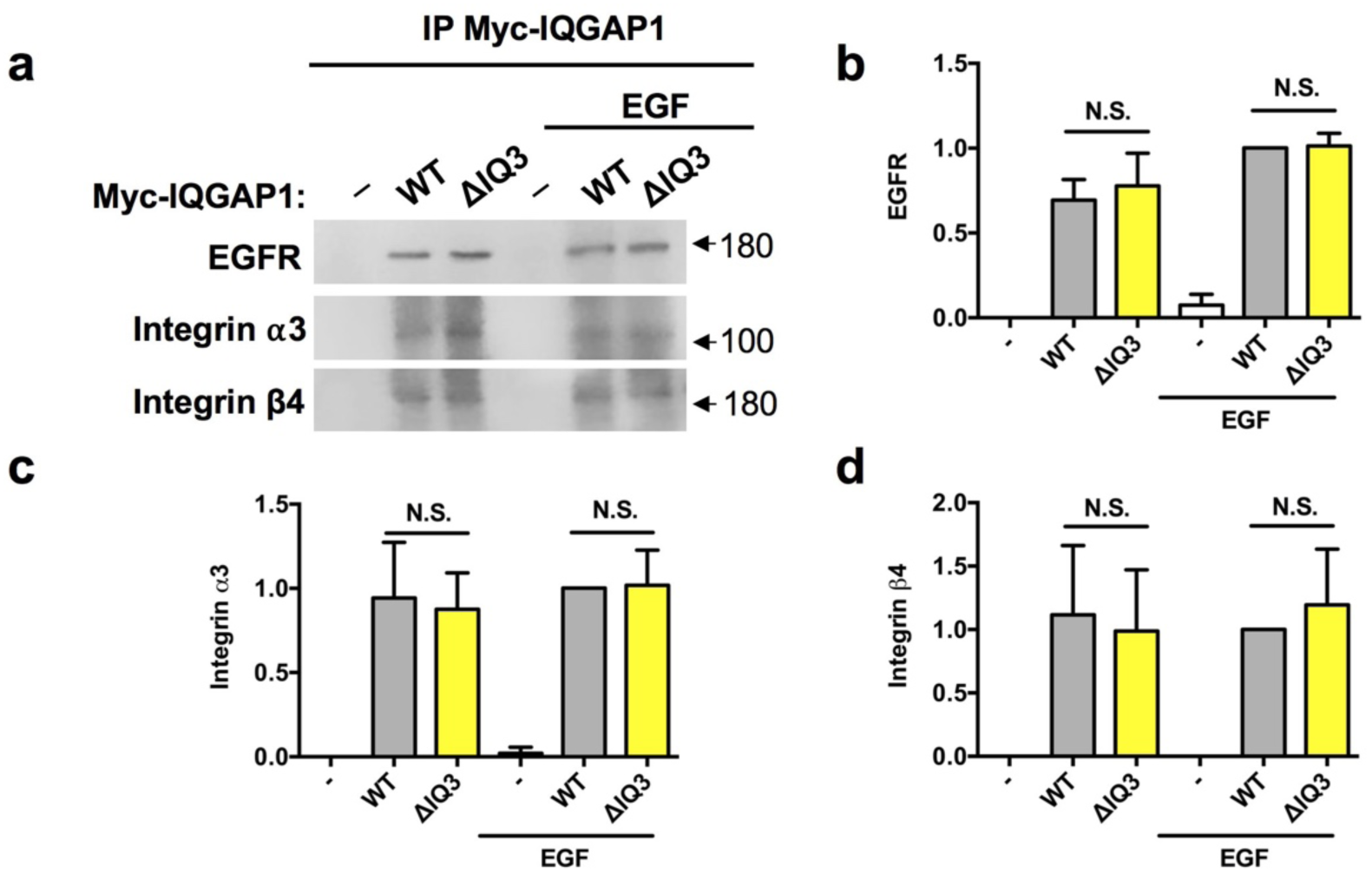
IQ3 deletion in IQGAP1 retains the binding with cell surface receptors. (**a**) IQGAP1 WT and ΔIQ3 mutant associated cell surface receptors were analyzed by immunoprecipitation. IQGAP1 WT and ΔIQ3 mutant interact with EGFR, Integrin α3 and Integrin β4 at similar levels. HEK293FT cells post 24 h of transient transfection with Myc-tagged IQGAP1^WT^ or Myc-tagged IQGAP1^ΔIQ3^ were starved for 24 h and then treated with 10 ng/ml EGF for 15 min as indicated. The whole cell lysates were collected and processed for immunoprecipitation using anti-Myc antibody conjugated Protein A/G agarose beads. (**b-d**) Quantification of immunoblots in **a**. n=3. Error bars denote SD.

To confirm this in cells, the interaction between WT/IQ3 deletion mutant of IQGAP1 and these cell surface receptors was determined by proximity ligation assay (PLA), a method allowing visualization and quantification of specific protein interaction events *insitu*^19^. The red fluorescent PLA signal represents the direct interaction or very close proximity between two proteins in their native state and cellular location. For this, HEK293FT cells transfected with Myc-tagged WT and IQ3 deletion mutant of IQGAP1 were starved and then stimulated by EGF before processing for PLA. Both WT and the IQ3 deletion mutant of IQGAP1 showed strong PLA signal with EGFR, integrin a3β1 and integrin a6β4 with or without EGF stimulation, consistent with an interaction with these cell surface receptors (Figure 6).

**Figure 6.**
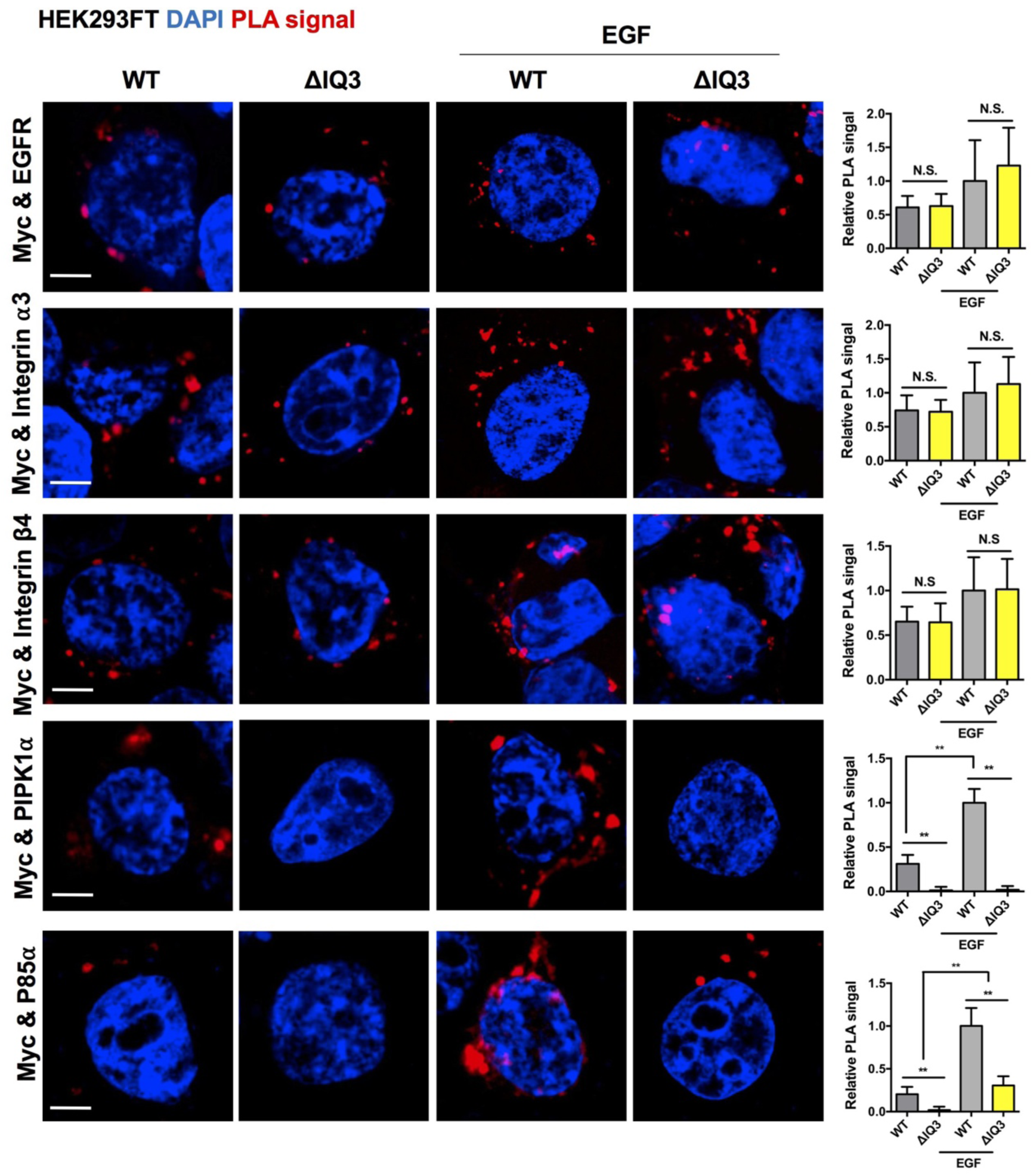
IQ3 mediates the binding of IQGAP1 with PI3K-Akt pathway components PIPK1α and p85. but not cell surface receptors in HEK293FT cells. PLA demonstrated the interaction between IQGAP1 WT/ ΔIQ3 mutant and EGFR/Integrin α3β1/Integrin α6β4/PIPK1α/p85α. HEK293FT cells post 24 h of transient transfection with Myctagged IQGAP1^WT^ or Myc-tagged IQGAP1I^ΔIQ3^ were starved for 24 h and then treated with 10 ng/ml EGF for 15 min. The cells were fixed and processed for PLA to determine the direct interaction between the Myc-tagged IQGAP1 WT/ΔIQ3 mutant and EGFR/Integrin α3/Integrin β4/PIPK1α/p85α. ⋆⋆P<0.01, n=10. Error bars denote SD. Scale bar, 5 μm.

We also examined the interaction between WT/IQ3 deletion mutant of IQGAP1 with the PI3K-Akt pathway component PIPK1α/p85α through PLA in HEK293FT cells. While the WT IQGAP1 and PIPK1α possessed a significant inducible PLA signal upon EGF stimulation, the IQ3 deletion mutant of IQGAP1 completely lost the PLA signal regardless of the EGF treatment (Figure 6). Similarly, the PLA signal between WT IQGAP1 and p85α was detectable under the starved condition and was also enhanced by EGF-stimulation. The PLA signal between IQ3 deletion mutant of IQGAP1 and p85α was partially reduced (Figure 6). In addition, there was no detectable PLA signal in the non-transfected control cells, confirming the validity of the interaction between the WT and IQ3 deletion mutant of IQGAP1 and cell surface receptors (Data not shown).

These experiments were replicated in the HNC cell line UM-SCC47. In these cells, the WT IQGAP1 possessed an inducible PLA signal with cell surface receptor EGFR and the PI3K-Akt pathway component PIPK1α upon EGF treatment (Figure 7). The IQ3 deletion mutant was able to sustain the PLA signal with EGFR at a comparable level to the WT IQGAP1 but lost the PLA signal with PIPK1α. Therefore, IQ3 deletion mutant retains the binding of IQGAP1 to cell surface receptors, whereas binding with the PI3K-Akt pathway components is compromised.

**Figure 7.**
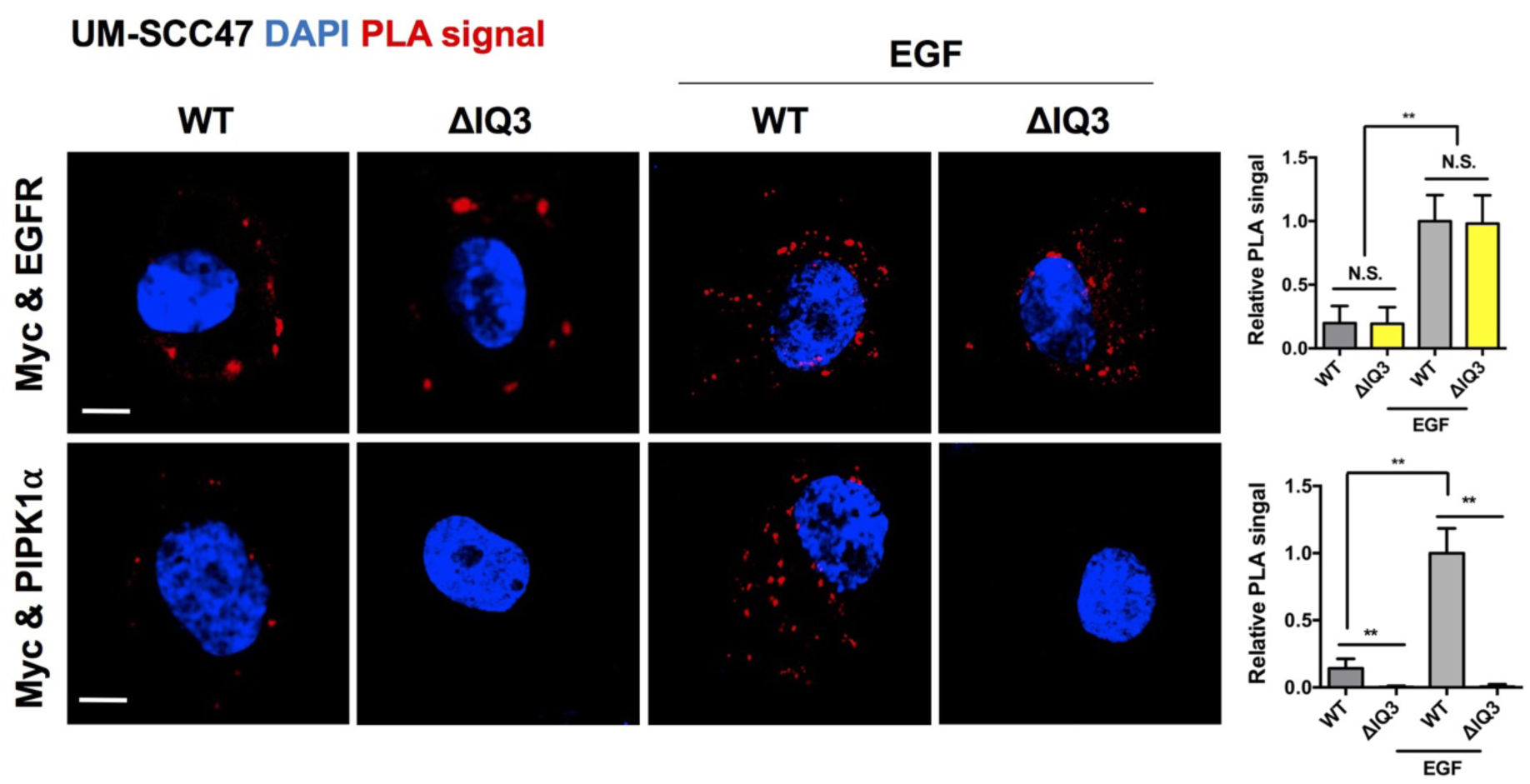
IQ3 deletion in IQGAP1 retains its interaction with EGFR but lost the interaction with PIPK1. in UM-SCC47 cells. PLA demonstrated interactions between IQGAP1 with EGFR and with PIPKIα. UM-SCC47 cells 24 h after transfection with Myc-tagged IQGAP1^WT^ or Myc-tagged IQGAP1^ΔIQ3^ were starved for 24 h and then treated with 10 ng/ml EGF for 15 min. The cells were fixed and processed for PLA to determine the direct interaction between the Myc-tagged IQGAP1 or IQGAP1^ΔIQ3^ and EGFR or PIPK1αα ⋆⋆P<0.01, n=10. Error bars denote SD. Scale bar, 5 μm.

### IQ3 motif mediates the role of IQGAP1 in cell migration

As IQGAP1 is a key regulator of cell adhesion and migration^20^, we examined the role of the IQ3 motif in the inducible function of IQGAP1 in cell migration the wound healing assay. The human breast cancer cell line MDA-MB-231 cells and the HNC cell line UM-SCC47 cells transfected with WT and IQ3 deletion mutant of IQGAP1 were cultured to confluency. These cells were starved and stimulated with EGF and the confluent cell layer in the plate was scratched using a plastic pipette tip. The migration of the cells at the edge of the scratch was imaged and analyzed at 5-time points up to 48 h (Figure 8). The results show that, in both MDA-MB-231 cells and UM-SCC47 cells, the WT IQGAP1 further promoted the EGF-stimulated wound healing process while the IQ3 deletion mutant of IQGAP1 lost this effect, demonstrating that the IQ3 motif is required for the inducible function of IQGAP1 in cell migration.

**Figure 8.**
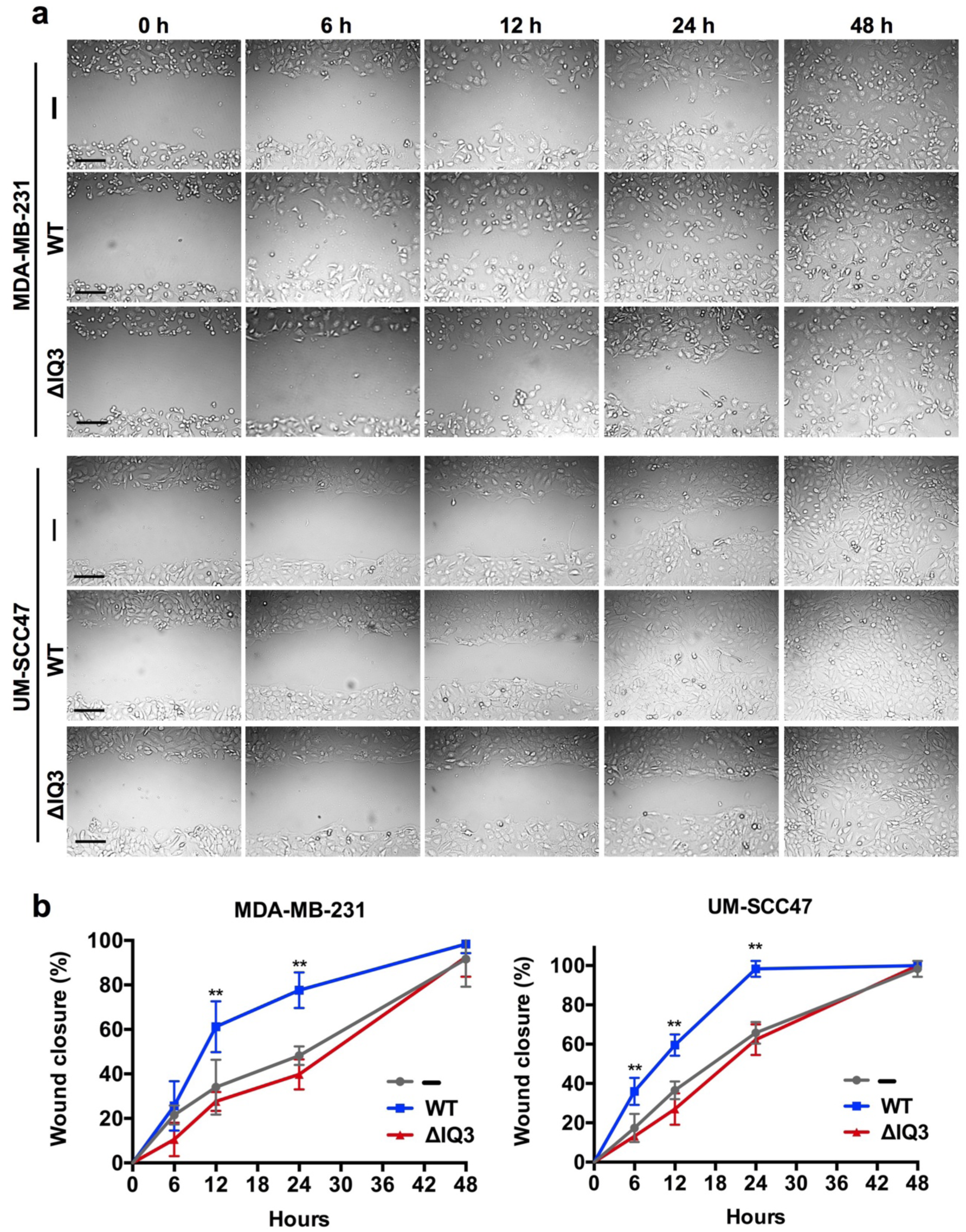
IQ3 motif regulates the inducible function of IQGAP1 in cell migration. (**a**) The MDA-MB-231 cells or UM-SCC47 cells were seeded and transfected with IQGAP1^WT^ or IQGAP1^ΔIQ3^ constructs for 24 h and then grown to confluence. The cells were starved in serumfree medium for 24 h and then treated with 10 ng/ml EGF. The cellular layer in each plate was scratched using a plastic pipette tip. The migration of the cells at the edge of the scratch was imaged at 0, 6, 12, 24 and 48 h. Scale bar, 100 μm. (**b**) Quantification of the percentage of wound closure at each time point. ⋆⋆*P*<0.01, n=3. Error bars denote SD.

### IQ3 motif-derived peptide diminishes the Akt activation and leads to the blockage of cell invasion

In cancer cells, IQGAP1 has a pivotal role in tumorigenesis and metastasis^21^. The agonist-stimulated scaffolding role of the IQ3 motif in IQGAP1 is likely to extend to the cell invasion. It has been reported that EGF-stimulated migration of epithelial cells depends on its incorporation into a receptor complex organized by syndecan-4 (Sdc4), in which EGFR and the laminin-binding integrin a3β1 engage a docking site in the extracellular domain of the syndecan, while the laminin-binding integrin a6β4 is engaged by the syndecan cytoplasmic domain (the Sdc4:EGFR:ITG complex)
^15,17^. The formation of this receptor complex and its ability to activate cell migration in response to EGF is blocked by a peptide mimetic of the docking site in the syndecan called SSTN_EGFR_^17^. Previously, we have demonstrated that a cell-penetrating IQ3 motif-derived peptide IQ3 is able to block the PI3K-Akt pathway^4^. To determine whether the IQ3 motif is specifically responsible for the scaffolding role of IQGAP1 at the downstream of the cell surface receptor stimulation, we assayed the Akt activation in UM-SCC47 cells treated with SSTN_EGFR_ and/or IQ3 peptides.

We found that Akt activation in UM-SCC47 cells was strongly reduced by treatment with 30 μM SSTN_EGFR_, or by treatment with 30 μM IQ3, whereas the two peptides together did not enhance inhibition (Figure 9a), suggesting that IQGAP1 and the Sdc4:EGFR:ITG complex are in the same pathway. To investigate the role of these signaling complexes in UM-SCC47 cell invasion through laminin-5 (LN-5, also called LN332)-coated filters (Figure 9b). As shown in Figure 9c, SSTN_EGFR_ effectively inhibited EGF-induced invasion, as did the EGFR kinase inhibitors gefitinib and erlotinib. Inhibition was also observed using the Src-family kinase PP2, an inhibition that is predicted based on the known role of Fyn in phosphorylating the integrin β4 subunit downstream of EGFR^22^. Invasion is also effectively blocked by the IQ3 peptide, as well as inhibiting PIPK1α activity with ISA2011B. Taken together, these results indicate that the IQ3 motif mediates PI3K and Akt activation downstream of the Sdc4:EGFR:ITG complex that mediates the invasion of these cancer cells.

**Figure 9.**
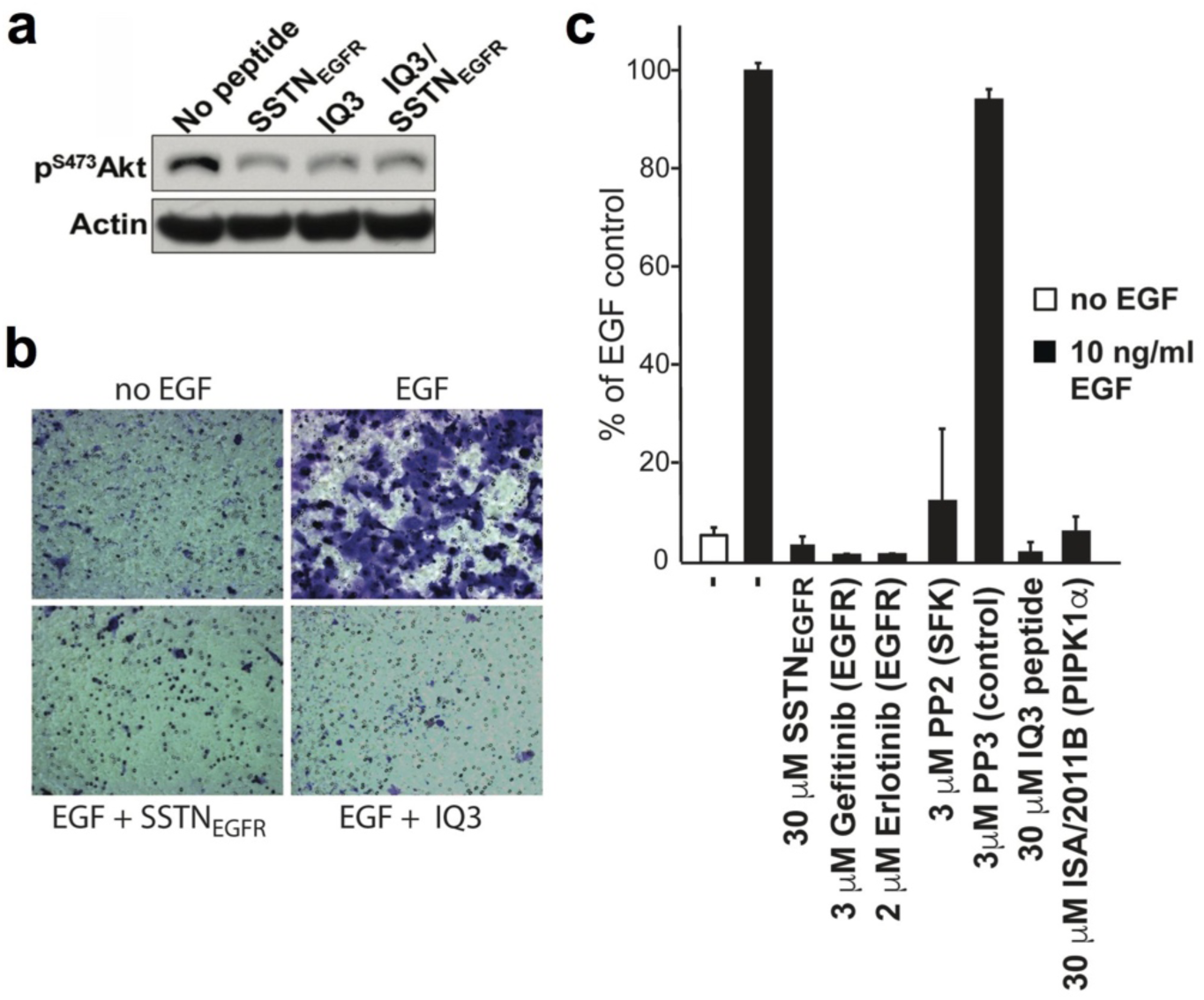
IQ3 motif mediates PI3K and Akt activation downstream of the Sdc4:EGFR:ITG complex that mediates invasion of UM-SCC47 cells. (**a**) Treatment of UM-SCC47 cells with 30 μM SSTN_EGFR_ or IQ3 peptide or both inhibited Akt activation. (**b**) Transfilter invasion of UM-SCC47 cells induced by EGF on LN5 is inhibited by 30 uM SSTN_EGFR_ or 30 μM IQ3 peptide. (**c**) Quantification of EGF-induced transfilter invasion in the presence of 30 μM SSTN_EGFR_, EGFR inhibitors (3 μM gefitinib, 2 μM erlotinib), 3 μM Src-family kinase inhibitor PP2 or its inactive analog PP3, 30 μM IQ3 peptide, or 30 μM PIPK1α inhibitor ISA/2011B. n=5. Error bars denote SD.

## Discussion

Here we have shown that IQ3 domain of IQGAP1 is a specific sequence required for the scaffolding of PI3K-Akt pathway components PIPKIα and p85α within the context of full-length IQGAP1 (Figure 10). Deletion of the IQ3 domain in IQGAP1 specifically blocked the agonist-stimulated PI3K-Akt pathway without impact on the IQGAP1 scaffolded Ras-ERK pathway. These findings have defined the function and specificity of IQGAP1 in PI3K-Akt pathway more precisely, leading towards the spatial and temporal regulation of the scaffolding machinery. These data indicate that the PI3K-Akt pathway, the Ras-ERK pathway and the interaction with cell surface receptors require distinct interaction sites on IQGAP1. The dissection of these interaction sites will allow the definition of the underlying mechanisms for IQGAP1 activation of each pathway.

**Figure 10.**
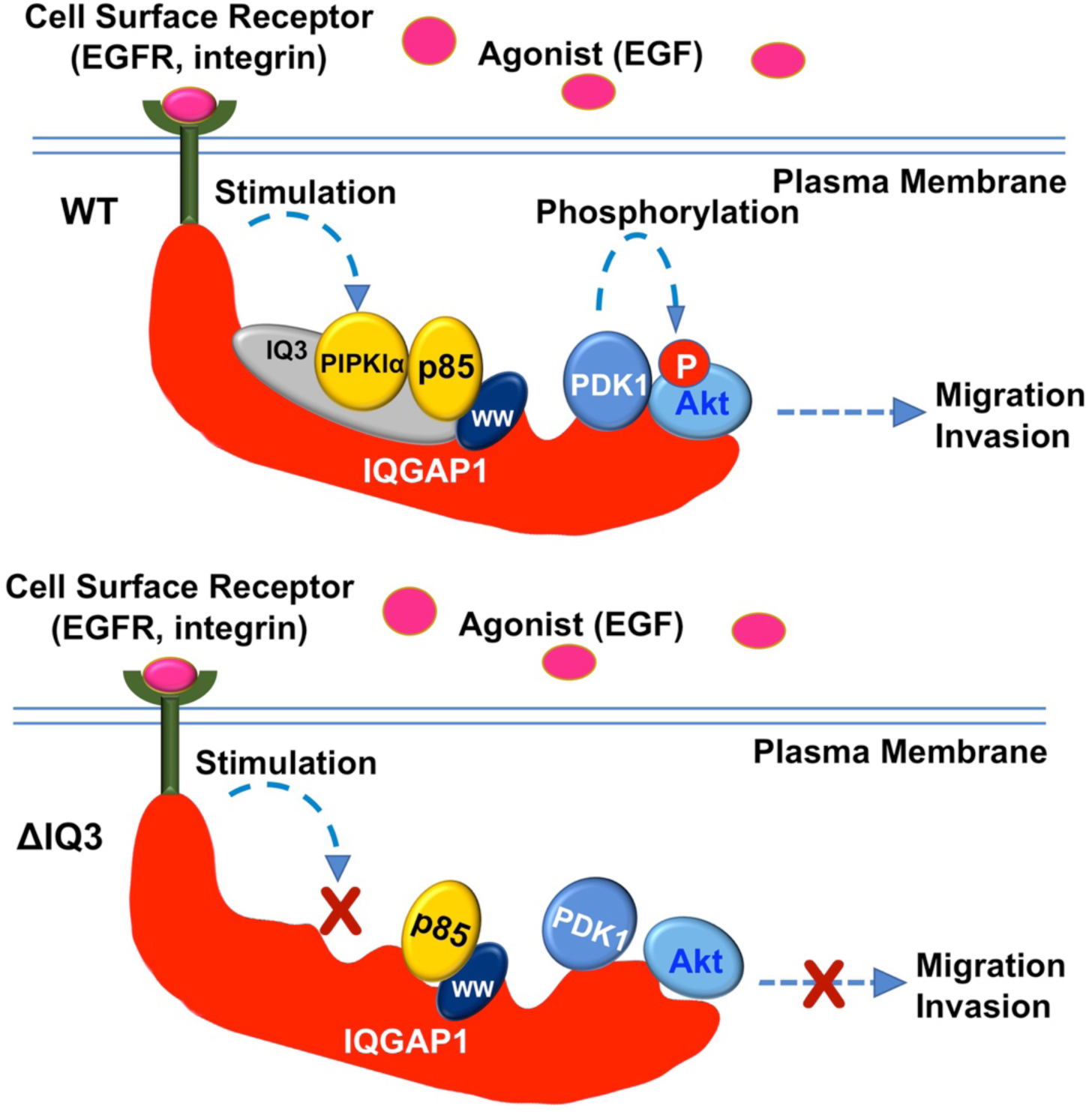
Proposed model for the role of the IQ3 motif in the IQGAP1 scaffolding of the PI3K pathway upon agonist stimulation.

IQGAP1 plays a crucial role in cell growth and survival^21^. Cancer cells with overexpressed IQGAP1 experience enhanced Akt activation^4^ and diminished ERK activation^23^. Our previous data suggested that many breast cancer cells become addicted to the IQGAP1-PI3K pathway for survival^4^. However, some cancer cells do not fully rely on one pathway. If the PI3K-Akt pathway is inhibited, in some cancers the Ras-ERK pathway is up regulated^4,12^, as phospho-FOXO1 that inhibits the Ras-ERK pathway is lost. By determining the specific binding site and regulatory mechanism of each pathway scaffolded by IQGAP1, we may be able to block the shifting of IQGAP1 scaffolded pathways and develop more efficient cancer therapies.

The IQGAP1 IQ3 deletion mutant lacks 20 amino acids in the IQ3 motif that is a shared binding region for PIPKIα and p85α. Future directions should include the definition of the exact binding sites of PIPKIα and p85α within IQ3 domain through making a series of point mutations. The candidate mutation sites will be determined by examining the conserved motifs or amino acid residues among different species in IQGAP1 or among members of IQGAP family that suggests crucial amino acids required for the PI3K-Akt pathway components scaffolding. Each pattern of point mutations will be assessed whether it resembles the effect of IQ3 deletion on the PI3K-Akt and Ras-ERK pathway. With the knowledge of binding sites of individual kinases, we can further investigate the mechanism of signaling and assembling of PI3K-Akt pathway complex by IQGAP1, and be able to design therapies to specifically eliminate one of the pathways IQGAP1 scaffolds in cancer cells.

IQGAP1 is known to play a pivotal role in the control of cell survival, proliferation, adhesion, polarization, membrane trafficking and invasion^20^. By specifically dissecting the scaffolding roles of IQGAP1 we now have the potential to define which scaffolded pathways function in specific IQGAP1 roles within cells. This, in turn, may allow for the design of target therapeutics that block specific signaling pathways and inhibit upregulation of others that may lead to therapeutic resistance. In addition, at IQGAP2 and IQGAP3 also function to scaffold signaling pathways these results will lead to an understanding of how these scaffolds control signaling.

In summary, this work has demonstrated the IQ3 motif as a specific target in IQGAP1 for the PI3K-AKT pathway. This region is selectively responsible for the scaffolding role of IQGAP1 to assemble PIPK1α and PI3K, key components of the PI3K-Akt pathway. This discovery opens the door for a novel therapeutic strategy targeting the PI3K-Akt signaling in cancers.

## Materials and Methods

### Cell culture and reagents

HEK293FT, MDA-MB-231, and UM-SCC47 cells were purchased from ATCC. HEK293FT and MDA-MB-231 cells were maintained in DMEM supplemented with 10% fetal bovine serum (FBS) (Gibco). UM-SCC47 cells were cultured in DMEM supplemented with 10% FBS and 1 mg/ml hydrocortisone (Sigma-Aldrich). Antibodies for immunoblotting, immunofluorescent (IF) staining and immunoprecipitation were from Cell Signaling Technology, including, pAkt (Ser473, 4058), ERK (4695), pErk (Thr202/Tyr204, 4370), PDK1 (13037), p85α (4292), PIPK1α (9693), Santa Cruz Biotechnology, including IQGAP1(H-109) and EGFR (SC-03), Abcam, including Akt (ab126811), Merck Millipore, Myc-tag (05-724), Sigma-Aldrich, including Myc-tag (C3956), Novus Biologicals, including integrin α3 (NBP2-48514) and R&D Systems, including integrin β4 (MAB4060). Secondary antibodies were from Thermo Fisher Scientific for IF and Santa Cruz Biotechnology for WB. Other chemicals were from Sigma-Aldrich. Oligo nucleotides were designed by SnapGene and ordered from Thermo Fisher Scientific. The SSTN_EGFR_ peptide (NHIPERAGSGSQVPTEPKKLEENEVIPKRISPVEESEDVSNKVSM)^17^ and the membrane penetrating IQ3 peptide ((dR)(dR)(dR)(dR)(dR)(dR)(dR)(dR)EQKLISEEDLEVVKIQSLARMHQARKRYRD)^4^ as previously described were synthesized by Genscript.

## Constructs

The Myc-tagged and GFP-tagged WT IQGAP1 (IQGAP1^WT^) constructs used for this work have been described previously^4,13^. Both Myc-tagged and GFP-tagged IQ3 motif deleted construct of IQGAP1 (IQGAP1^ΔIQ3^) were generated by the overlapping polymerase chain reactions (PCR) to delete the 20 amino acids (EVVKIQSLARMHQARKRYRD) in the IQ3 motif. The IQ3 motif in IQGAP1 located within two XcmI enzyme sites. The sense primer 5’-CTGCGCTCCCACAAAGATCGCCTGCAGTACTTCCGG-3’ (forward linker) and anti-sense primer 5’-CCGGAAGTACTGCAGGCGATCTTTGTGGGAGCGCAG-3’ (reverse linker) were used to link the flanking region of the deleted IQ3 motif. The DNA fragment at the upstream of the IQ3 motif was amplified using the primer with XcmI cutting site 5’-GAGTCCATGAACTTGGTGGACTTG-3’ (XcmI forward) and the reverse linker. The DNA fragment at the downstream of the IQ3 motif was amplified using the forward linker and the primer with XcmI cutting site 5’-CCTCCCACCACAGAACTGGAGGA-3’ (XcmI reverse). The upstream and downstream PCR products of IQ3 region was joined based on their overlapping linker, amplified using above mentioned XcmI forward and XcmI reverse primers and cloned in the Myc-tagged and GFP-tagged IQGAP1^WT^ constructs within the XcmI enzyme site to replace the WT IQGAP1 into the ΔIQ3 deletion mutant.

## Transfection

Constructs were transfected into cells by a lipid-based delivery system from Thermo Fisher Scientific. The cells were cultured to be 70% confluent at transfection. For 1×10^6^ cells, 5 μL of Lipofectamine^TM^ 3000 reagent was diluted in 250 μL Opti-MEM medium. Two μg of indicated IQGAP1 plasmid was added into 250 μL Opti-MEM medium containing 5 μL of P3000 reagent. The diluted lipofectamine reagent and diluted DNA were mixed and incubated at room temperature for 10 min. Then the DNA-lipid complex was added to the cells drop wisely. The cells post 48 h transfection were ready for visualization and further analysis.

## Immunofluorescent staining

HEK293FT and MDA-MB-231 cells were seeded onto 0.2% Gelatin pre-coated glass coverslips of thickness No. 1.5H. Cells post 24 h transient transfection of GFP-tagged IQGAP1^WT^ or GFP-tagged IQGAP1^ΔIQ3^ were starved for 24 h in serum-free medium and then stimulated with 10 ng/ml EGF for 15 min. Then the cells were fixed by 4% paraformaldehyde (PFA) in phosphate-buffered saline (PBS) for 30 min in room temperature. After 3 times wash by PBS, the cells were permeabilized with 0.3% Triton in PBS for 15 min. The cells were washed again for 3 times by PBS and blocked with 3% Bovine Serum Albumin (BSA) in PBS for 1 h at room temperature. After blocking, cells were incubated with primary antibody at 1:300 dilution in PBS containing 3% BSA at 4°C overnight. After washing 3 times by PBS with 0.1% Tween20 (PBST), cells were stained with corresponding secondary antibody for 1 h at room temperature. Cells were again washed 3 times with PBST before being mounted with Fluoroshield mounting buffer with DAPI.

## Confocal microscopy

Images of fixed cells were collected by Leica SP8 confocal microscope, which includes a super-supercontinuum white-light laser for fluorescent excitation from 470nm to 670nm, a separate diode laser at 405nm. This microscope is equipped with 3 Photomultiplier Tubes (PMTs) and 2 high-sensitive HyD detectors for image collection. All confocal images were acquired using the 100x objective lens (N.A. 1.4 oil). This microscope was controlled by Leica LAS software. The fluorescent intensity of each channel was measured by ImageJ. The quantitative graph was generated by GraphPad Prism version 6.0 software.

## Immunoblotting and immunoprecipitation

HEK293FT cells were transiently transfected with Myc-tagged IQGAP1^WT^ or Myc-tagged IQGAP1^ΔIQ3^ construct. After 24h, the cells were starved for 24 h in serum-free medium and then stimulated with 10 ng/ml EGF for 15 min. Cells were lysed in a buffer containing 25 mM Tris-HCl pH 7.4, 150 mM NaCl, 1 mM EDTA, 1% NP-40, 5% glycerol, 5 mM Na_3_VO_4_, 20 mM NaF and a protease inhibitor cocktail from Roche. The protein concentration of lysates was measured by Bradford Protein Assay (Bio-Rad) and an equal amount of protein was used for further analysis. All primary antibodies were at 1:1000 dilution for immunoblotting. For immunoprecipitation, 0.5 mg of proteins were incubated with 20 μL of anti-c-Myc antibodies pre-bound agarose for 2 h at 4°C. After incubation, the agarose beads were washed 3 times with lysis buffer and eluted with SDS sample buffer. For WB, 10 μg of proteins were loaded for each lane. Protein bands were quantified by ImageJ and the statistical analysis was performed with GraphPad Prism version 6.0 software. The statistical analysis was generated using data from three independent experiments.

## Proximity ligation assay

HEK293FT cells or UM-SCC47 cells were seeded to 0.2% Gelatin-coated glass cover slides and transiently transfected with Myc-tagged IQGAP1^WT^ or Myc-tagged IQGAP1^ΔIQ3^ construct. Then the cells were starved for 24 h in serum-free medium and stimulated by 10 ng/ml EGF for 15 min. After that, the cells were fixed by 4% PFA and processed for proximity ligation assay (PLA). The interaction between the Myc-tagged WT/ΔIQ3 of IQGAP1 and EGFR/Integrin α3/Integrin β4 were determined by the red PLA signals from Duolink *in situ* red starter kit mouse/rabbit (Sigma-Aldrich). The untransfected cells without Myc-tagged IQGAP1 expression were used as the control. The images were collected by Leica SP8 confocal microscope and analyzed by ImageJ. The PLA signal in 10 cells of each group was used for the quantification.

## Wound healing assay

The MDA-MB-231 or UM-SCC47 cells (1×10^5^ cells/35×11 mm dishes) were seeded and transfected by IQGAP1^WT^ or IQGAP1^ΔIQ3^ constructs for 24 h before achieving confluence. Then the cells were starved in serum-free medium for 24 h and treated with 10 ng/ml EGF. The cellular layer in each plate was scratched using a plastic pipette tip. The migration of the cells at the edge of the scratch was analyzed at 0, 6, 12, 24 and 48 h, when microscopic images of the cells were captured using a 10x objective on a Nikon Eclipse TE2000U equipped with a Photometrics coolSNAP ES CCD camera. The system was controlled by MetaMorph v6.3 (Molecular Devices). Three random fields were imaged and later quantified by ImageJ.

## Invasion assay

The bottom chambers of transwell filter chambers (8 μm pores; Corning) were coated with 10 μg/ml of LN-5 in PBS for 3 h. UM-SCC47 cells (5 x 10^4^) suspended in DMEM containing 1% BSA (serum-free) were placed in the upper chamber with or without indicated inhibitors and incubated for 16 h at 37°C. Cells on the bottom of the filter were fixed with 4% Paraformaldehyde diluted in PBS and stained with 0.1% Crystal Violet and five random fields were imaged and counted.

## Statistical analysis

Data were expressed as the mean ± standard deviation (±SD). Statistical significance was determined using student t-test. *P<0.05; **P<0.01, n ≥3.

## Acknowledgements

We would like to thank Lance Rodenkirch from the Optical Imaging Core of UW-Madison for the technical support of confocal microscopy and all members of the R.A.A., A.C.R., and P.F.L. groups. This work was supported by National Institutes of Health (NIH) grants to R.A.A. (R01-GM57549, R01-CA104708), A.C.R. (R01-CA163662) and P.F.L. (R35-CA 210807, P01-CA022443), support to A.C.R. and P.F.L. from the UW Head and Neck Cancer SPORE (P50DE026787), funds to support Head and Neck cancer research from the UW School of Medicine and Public Health and the UW Carbone Cancer Center to A.C.R, P.F.L., and R.A.A., and the use of the cancer center shared services, supported by NIH/NCI P30 CA014520.

## Authorship Contributions

M.C. participated in concept design, generation of data and writing the manuscript; T.W. participated in writing the manuscript and figure preparation; S.C. and O.J. performed the experiments; N.T. provided critique of the manuscript; A.C.R. and P.F.L. participated in concept design and critique of the manuscript, and R.A.A. supervised concept design and participated in writing the manuscript.

